# Determination of expression profiles for Drosophila ovarian Follicle Stem Cells (FSCs) using single-cell RNA sequencing

**DOI:** 10.1101/2022.06.27.497544

**Authors:** Zhi Dong, Lan Pang, Zhiguo Liu, Yifeng Sheng, Xavier Thibault, Amy Reilein, Daniel Kalderon, Jianhua Huang

## Abstract

Drosophila ovarian Follicle Stem Cells (FSC) present an excellent paradigm for understanding how a community of active stem cells maintained by population asymmetry is regulated. Here we describe single-cell RNA sequencing studies of a pre-sorted population of cells that include FSCs and the neighboring cell types, Escort Cells (ECs) and Follicle Cells (FCs), which they support. Cell-type assignment relies on anterior-posterior (AP) location within the germarium. We clarify the previously determined location of FSCs and use spatially targeted lineage studies as further confirmation. The scRNA profiles among four clusters are consistent with an AP progression from anterior ECs through posterior ECs and then FSCs, to early FCs. Several genes with graded profiles from ECs to FCs are highlighted as candidate effectors of the inverse gradients of the two principal signaling pathways, Wnt and JAK-STAT, that guide FSC differentiation and division.

## Introduction

Adult stem cells provide a lifelong source of specific differentiated cells, necessitated by physiological turnover or changing environmental conditions (1-3). The study of adult stem cells is revealing that this task can be accomplished in a variety of ways. Importantly, it is often a shared task. In such paradigms, often termed population asymmetry, individual stem cells make stochastic decisions about cell division and differentiation (2-8). Division and differentiation are independent processes in the three best-studied paradigms (9-11) and are balanced at the community level. The stochastic variation in behavior of individual stem cells may be compounded further by systematic heterogeneity based on precise stem cell location (2, 12). For example, within a population of about 16 Follicle Stem Cells (FSCs) in a Drosophila germarium, roughly half are in a posterior ring. Only posterior FSCs can directly become proliferative Follicle Cells (FCs) and they also divide much faster than their anterior FSC neighbors, which can directly become quiescent Escort Cells (ECs) (13). Posterior and anterior FSCs nevertheless form a single community because they can exchange positions (13). It has also often been observed that stem cell derivatives can revert to stem cell status under physiological or stress conditions (14-16). All of these plastic properties, including variable stem cell lifetimes, differ substantially from the original concept, still relevant to some paradigms, of each stem cell behaving the same way-dividing with irreversible asymmetric outcomes for the two daughters and exhibiting exceptional longevity (1-3, 7).

Many aspects of the status of a cell can be captured by a detailed rendition of its pattern of RNA expression, so understanding of adult stem cell biology surely benefits from such information, typically captured by single-cell RNA (scRNA) sequencing. However, adult stem cell paradigms that exhibit stochastic and heterogeneous behaviors, with derivative cells that are necessarily initially very similar and do not necessarily quickly transit to an irreversibly distinct state, present a significant challenge to categorization of stem cells and their immediate derivatives according to scRNA profiles. These profiles cannot be expected to reveal stem cells as a highly distinct group and certainly cannot suffice to define stem cells. Instead, they must be related carefully to the results of functional tests that define stem cells through their behavior (2). Drosophila FSCs present a particularly attractive paradigm for such analyses because their behavior is complex but has been studied carefully, along with extensive investigation of key regulatory niche signals (2, 9, 13, 17, 18). Here, we present a scRNA study of somatic ECs, FSCs and early FCs within the Drosophila germarium and we relate those studies to functional tests of FSC behavior based on location. The resulting expression profiles reveal candidate positionally-regulated effectors of FSC division and differentiation.

## Results

At the anterior of each germarium are non-dividing somatic terminal filament (TF) and cap cells, which contact 2-3 germline stem cells (GSCs) (Fig. 1A). GSCs generally divide asymmetrically to produce a more posterior Cystoblast, which divides four times with incomplete cytokinesis (19-21). The developing germline cyst derivatives are wrapped by processes of somatic ECs as they progress posteriorly through region 1 (22, 23) to form a rounded 16-cell stage 2a cyst, which then elongates into a lens-shaped 2b cyst, spanning the widest part of the germarium. Proliferative Follicle Cell precursors (“FCs”) associate with the 2b cyst, which then rounds to become a stage 3 cyst before budding from the germarium as an egg chamber, enclosed in a monolayer epithelium of FCs. Quiescent polar cells at the anterior and posterior termini of the FC epithelium contact non-dividing stalk cells, which connect consecutive egg chambers. Polar and stalk cell precursors are specified early within the germarium (24-26), while other FCs continue to divide until mid-oogenesis (stage 6), later specializing according to their AP and DV locations on the egg chamber (27, 28). FSCs lie between ECs and the earliest FCs, providing a continuous supply of new FCs and also, less frequently, replenishing ECs (13, 25).

**Figure 1.**
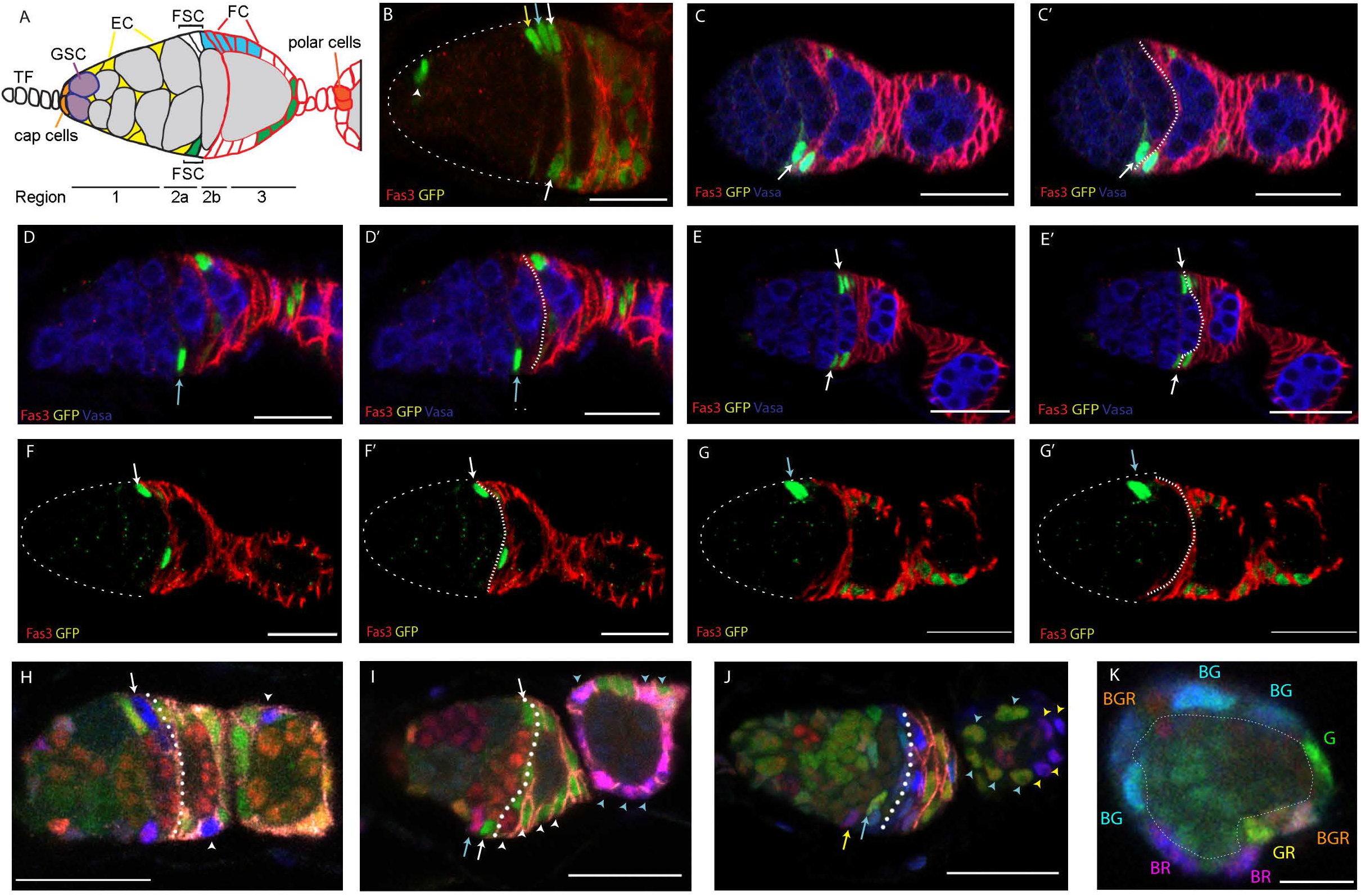
Location of FSCs in the Drosophila germarium. (A) Germline Stem Cell (GSC) daughters (gray) develop into 16-cell cysts as they progress further posterior (right) in region 1, supported by interactions with quiescent enveloping Escort Cells (ECs) and span the width of the germarium in region 2b as ECs are replaced by Follicle Cells (FCs). Two potential marked FSC lineages are shown (blue, green), together with the pattern of strong Fas3 surface protein expression (red). (B-J) Examples of FSC lineages marked by positive marking and multicolor lineage tracing. The anterior border of strong Fas3 generally aligns with the posterior of the stage 2b germline cyst, as in (A), and is indicated by broken white lines in (C’, D’, E’, F’, G’, H-J); central z-sections are shown but experiments examine cells in all z-sections. (B-G) show GFP-positive (green) MARCM FSC lineages with Fas3 (red) and (C-E) Vasa (blue) marking germline cells. FSC nuclei are indicated by colored arrows according to AP location of layer 1 (white), layer 2 (cyan) or layer 3 (yellow), from posterior to anterior. (B) Most FSC lineages that survive for several days contain several FSCs (arrows) and one or more labeled EC (arrowhead), precluding identification of which of these cells may be maintaining the lineage. Lineages with only a single candidate stem cell (C, D, F, G) or with candidates in only a single AP plane (E) were comprehensively scored to reveal that an FSC could be found immediately anterior to the Fas3 border (white arrows), one cell further anterior (blue arrows) or occasionally one cell further anterior. (H-K) Multicolor FSC lineages, marked by the loss of GFP (G, green), RFP (R, red) or lacZ (B, blue) in different combinations were analyzed in the same way to reveal single candidates or single-plane candidates, as indicated by colored arrows (derivative FCs of each are indicated by arrowheads of matching color). Single candidate FSCs were also found to be evenly distributed around the germarial circumference (in all z-sections). (K) A germarium oriented perpendicular to others shows that in cross-section around the AP plane of layer 1 FSCs there are multiple somatic cell nuclei (nine here, labeled according to retained colors) surrounding a central germline cyst. Scale bar 20 µm in (B-J), 10µm in (K).

Some scRNA studies have been conducted on whole ovarioles to capture all aspects of oogenesis (18, 29, 30). We were interested only in examining FSCs and their neighbors in detail, so we chose a strategy that began with purifying those cells. The enhancer trap line *C587-GAL4*, which is expressed in adult ECs, FSCs and the earliest FCs, was used to drive *UAS-CD8-RFP* cell surface protein expression, followed by manual dissection of ovaries, dissociation and fluorescence-activated cell sorting (FACS). Single-cell RNA sequences were derived on the 10XGenomics platform with unique molecular identifiers (UMIs). Average raw sequence reads per cell were over 290, 000. After genome alignment and UMI counting, genes featured in more than three cells were included, and cells with feature counts between 100 - 4,500, and fewer than 20% mitochondrial counts were retained for principal component analysis using Seurat (v4.0.2).

Following initial clustering of scRNA patterns for a total of over 1300 cells, we examined each group for known indicators of EC, FSC or early FC identity. We recognized small groups with features of cap cells or TF cells (selective expression of *Lmx1, en, dpp* and *wg*; 43 cells) (31-33), stalk cells and their precursors (selective expression of *LamC* and *sim*; 23 cells) (30), and germline cells (selective expression of *zpg, chinmo, ovo, otu* and *vasa*; 34 cells) (Fig. S1) (30). Three groups (0, 4, 9 in Fig. 2A) were not considered further on the basis of a combination of higher mitochondrial RNA content and lower total RNA counts, suggesting the possibility of damage or stress (Fig. 2B). The remaining four groups had characteristics of ECs and FSCs (*Wnt4, ptc, fax*) (13, 34-36) or early FCs (*Fas3, cas*) (24) and were re-sorted by themselves in order to gain better resolution among somatic cells of the anterior half of the germarium. The result was a progression of six clusters, roughly consistent with progressive anterior to posterior identities (Fig. 2C, D), as described further below.

**Figure 2.**
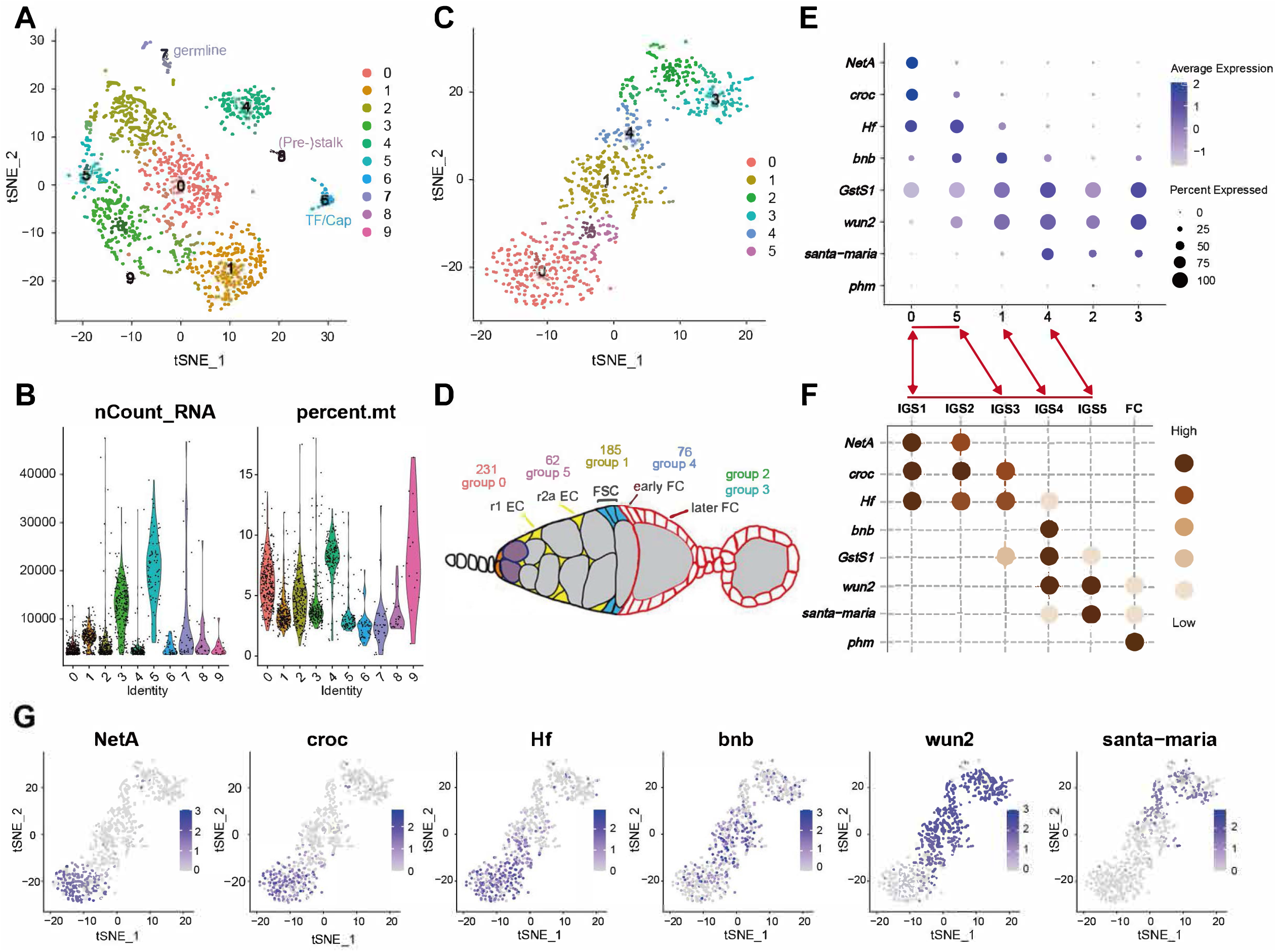
Cluster analysis of scRNA-seq profiles of germarial cells. (A) t-Distributed Stochastic Neighbor Embedding (t-SNE) plot of the dataset, together with (C) the number of unique molecular identifiers (nUMI) detected per cell and percentage of mitochondrial gene for each cluster. (B) t-SNE plot after re-clustering the original groups 1, 3, 5, 2 only. (D) Deduced location in the germarium of cells within the six clusters shown in (C). (E) Summary dot plot of key marker genes defining clusters presented in (37), together with dot plots of the same genes for the six clusters in (C). Double-headed arrows indicated similar groups in the two studies (groups 0 and 5 span the content of groups IGS1-3; *phm* was not detected in our study). (G) Expression of the six most informative maker genes imposed on t-SNE plot of (C) groups 0-5.

We compared our clusters to those derived from another study that also concentrated on anterior somatic germarial cells (37) and found substantial similarities. The patterns of prominent selectively expressed RNAs that defined groups previously labeled as “IGS1-5” (IGS, or inner germarial sheath, is an alternative name for Escort Cells) showed largely good correspondence to our group 0 (IGS1 and 2; *NetA* + *Croc*), 5 (IGS3; *Croc* + *Hf*), 1 (IGS4; *bnb* + *wun2*) and 4 (IGS5; *wun2* + *santa-maria*) (Fig. 2E-G). Importantly, Tu et al., (37) used several of these “marker RNAs” for *in situ* hybridization to define the spatial limits of specific clusters, confirming the anterior to posterior progression of groups 0-1-5-4 in that order, as described below. Clusters 2 and 3 appear to represent more mature FCs, with ribosomal protein RNAs dominating the most up-regulated genes of cluster 2 (Fig. 3E), consistent with the highly proliferative nature of FCs. This feature is presaged in the earliest FCs in group 4 by strong expression of two ribosomal protein RNAs (*RpL35* and *RpS6*) and *myc* (Fig. 3D). A handful of cells within group 2 expressed JAK-STAT ligands, *upd1* and *upd3*, characteristic of polar cells (38).

**Figure 3.**
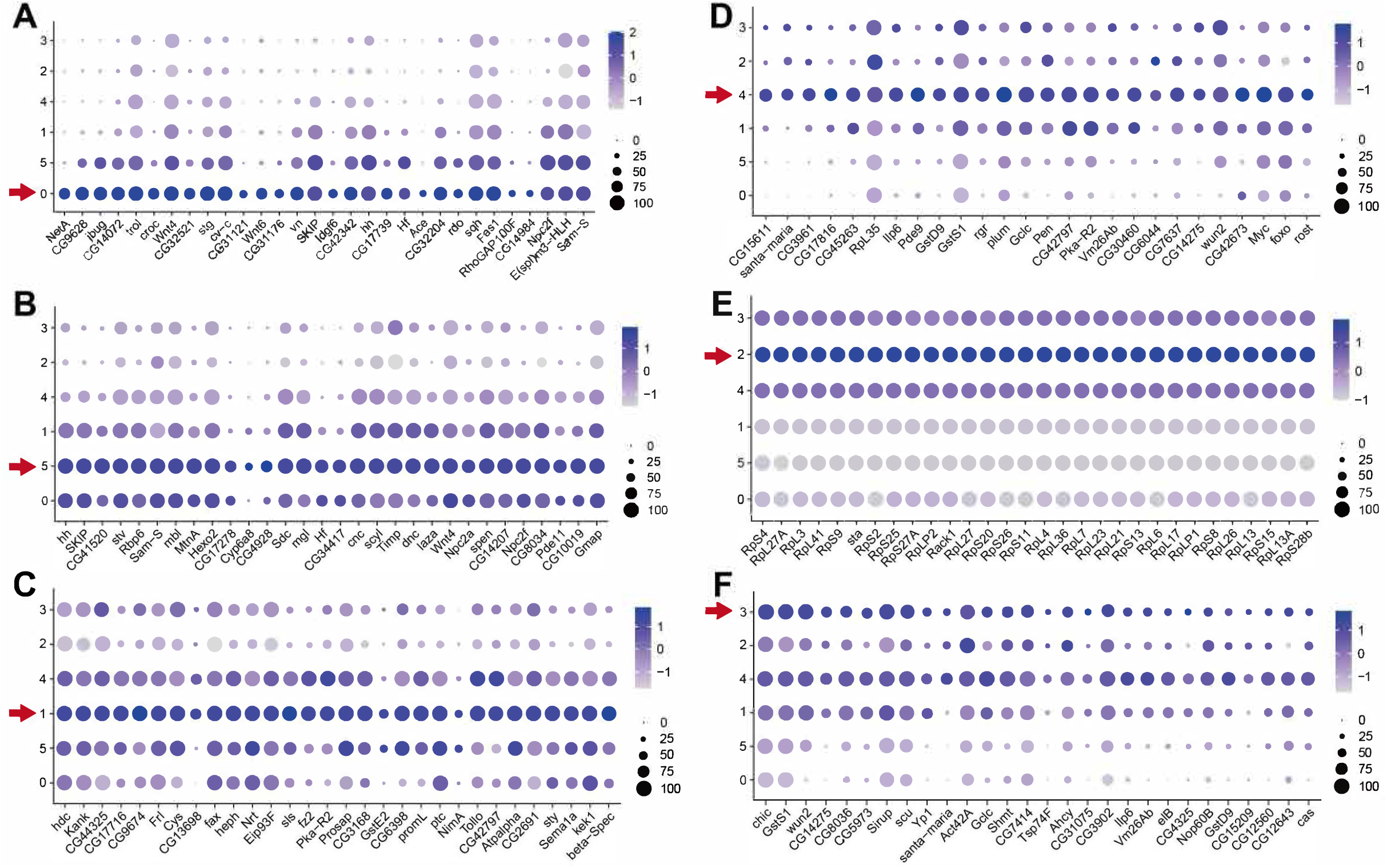
Marker genes for six clusters of somatic germarial cells. Dot plots showing the expression profiles of the top 30 markers (ordered by Adjusted p-value) in each of the clusters, (A) cluster 0, (B) cluster 5, (C) cluster 1, (D) cluster 4, (E) cluster 2 and (F) cluster 3, proceeding from anterior to posterior cell groups in the germarium.

### Cluster boundaries and cell identities

A landmark that has been used to define FSCs by functional assays is the anterior border of strong surface Fas3 staining (Fig. 1A). In key functional studies this landmark has been defined in a consistent manner (13, 17, 39). However, this is a challenging task to reproduce accurately over time with different operators because the three-dimensional structure of individual germaria can be irregular and a dynamic process is being sampled at random times throughout a 12h cycle of FC recruitment. We therefore describe a detailed protocol (see Methods) to promote consistency for all investigators.

The location of FSCs has previously been inferred by examining a large number of marked FSC lineages and using those with only a single candidate stem cell to determine FSC locations (2, 13). This strategy indicated that FSCs were almost all either immediately anterior to the strong Fas3 border (layer 1) or one cell further anterior (layer 2), with a few FSCs in the next most anterior location (layer 3). The total average number of layer 1 (eight) and layer 2 cells (six), distributed in radially equivalent locations abutting the germarial wall, was consistent with estimates of total FSC numbers derived from (i) measuring the average contribution of a single FSC lineage to all FCs and (ii) counting the number of different multicolored lineages in a single ovariole (in each case, taking steps to ensure capturing the full diversity of all FSC lineages, including the most short-lived) (2, 13). After defining FSC locations in this way, it was then possible to count labeled FSCs in layers 1 and 2 explicitly in numerous studies exploring the effects of altered genotypes, regardless of the number of marked FSCs per germarium (17). Some examples of layer 1 and 2 FSCs and their relationship to the strong Fas3 border are shown in Figure 1. Changes in the measured numbers and behaviors of marked FSCs due to altered genotypes in these locations formed a self-consistent picture of FSC responses to different signaling environments (17). Moreover, measurements of absolute rates of FSC division and conversion to FCs and ECs also matched (40), consistent with maintaining a stable population of about 16 FSCs, mainly in layers 1 and 2.

From RNA *in situs, santa maria* is mostly expressed posterior to the strong Fas3 border, with weaker expression just anterior to the Fas3 border, overlapping with the posterior margin of *bnb* RNA (37). *Santa maria* RNA is prevalent in group 4 but almost completely absent from group 1, while *bnb* 4 RNA is highest in group 1 but present in some cells of group 4, suggesting that the key Fas3 border is roughly between groups 1 and 4, with most FSCs in group 1 (Fig. 2F, G). A few of the most posterior FSCs, expressing both *bnb* and *santa-maria*, may be in group 4.

*bnb* expression was strongest in group 1 but also significant in abundance and prevalence in group 5 (Fig. 2F, G). Since the anterior boundary of *bnb* RNA expression *in situ* appears to extend beyond FSCs into r2a ECs (37), the transition between ECs and FSCs is likely very close to the border between groups 5 and 1. Other genes with sharp declines from group 5 to 1 include *Hf, CG34417* and *CG17278*, with the converse pattern for *CG13698, CG42797* and *fz2* (Fig. 3). Several RNAs peak within group 1 (*hdc, kank, fax, sls, promL, sty*), including *fax* (Fig. 3), for which a protein trap shows a maximum centered on the FSC domain (13).

Regarding the more anterior clusters, *NetA* RNA is restricted to within region 1, while the posterior border of *croc* RNA extends into region 2a, declining quite sharply at the anterior border of *bnb (37). NetA* expression was limited to group 0, while *croc* RNA was present also in a minority of group 5 cells, suggesting that groups 0 and 5 largely represent region 1 and 2a ECs, respectively (Fig. 2F, G).

Considering only groups 0, 5 and 1 (478 cells), the total number of cells in groups 0 (231, 48%), 5 (62, 13%) and 1 (185, 39%) are roughly in proportion to the average number of r1 Ecs (26, 46%), r2a (14, 25%) and FSCs (16, 29%) in an adult germarium (13, 41, 42), suggesting rough equivalence if all cells were captured equally well for sequencing. Thus, group 1 cells appear to correspond largely to FSCs, though it is expected that a few FSCs may also be within group 4 and not all cells in group 1 are necessarily FSCs (some r2a ECs and some FCs) (Fig. 2D).

### FSC gene expression patterns

One motivation for scRNA-seq studies is to identify candidate effectors of FSC behavior. The two basic behaviors of FSCs are division and differentiation. Functional studies have identified the strength of Wnt and JAK-STAT signaling pathways, with opposing intersecting gradients over the FSC domain, as major determinants of FSC position, differentiation to ECs at the anterior or FCs at the posterior (13, 17, 35). Consequently, it is expected that several downstream responders to these graded signals that execute differentiation responses will have an AP graded expression pattern around the FSC domain. Adhesion proteins are likely important effectors of these signals, potentially promoting posterior FSCs joining a nascent FC epithelium, anterior FSCs moving towards ECs or FSCs moving along a gradient of ECM adhesion ligands. We therefore looked for adhesion molecules (and some partners and regulators) with a clear gradient of expression from group 5 through group 1 to group 4 (Fig. 4A, B). Among the molecules with increasing expression from ECs (group 5) to FCs (group 4) were *Fas3, DE-cadherin* (*shg*), *Plum, Tetraspanin 74F* (*Tsp74F*), *bazooka* (*baz*), and *coracle* (*cora*) (*Dystroglycan* (*Dg*) and *Dystrophin* (*Dys*) plateau or peak in group 1). The opposite pattern was identified for *Fasciclin I* (*Fas1*), *echinoid* (*ed*), *Neuroglian* (*Nrg*), *Tenascin accessory* (*Ten-a*), *Tetraspanin* 66E (*Tsp66E*), *scab α-PS3 integrin* (*scb*), *Adherens junction protein p120* (*p120ctn*), *Zyxin* (*Zyx*) and *Lachesin* (*Lac*). Among these genes, only the role of DE-cadherin in FSCs has so far been investigated (43). A more comprehensive listing of expression in different clusters of genes with selectively strong expression in clusters 0, 5, 1 and 4 is presented in Supplementary Table 1.

**Figure 4.**
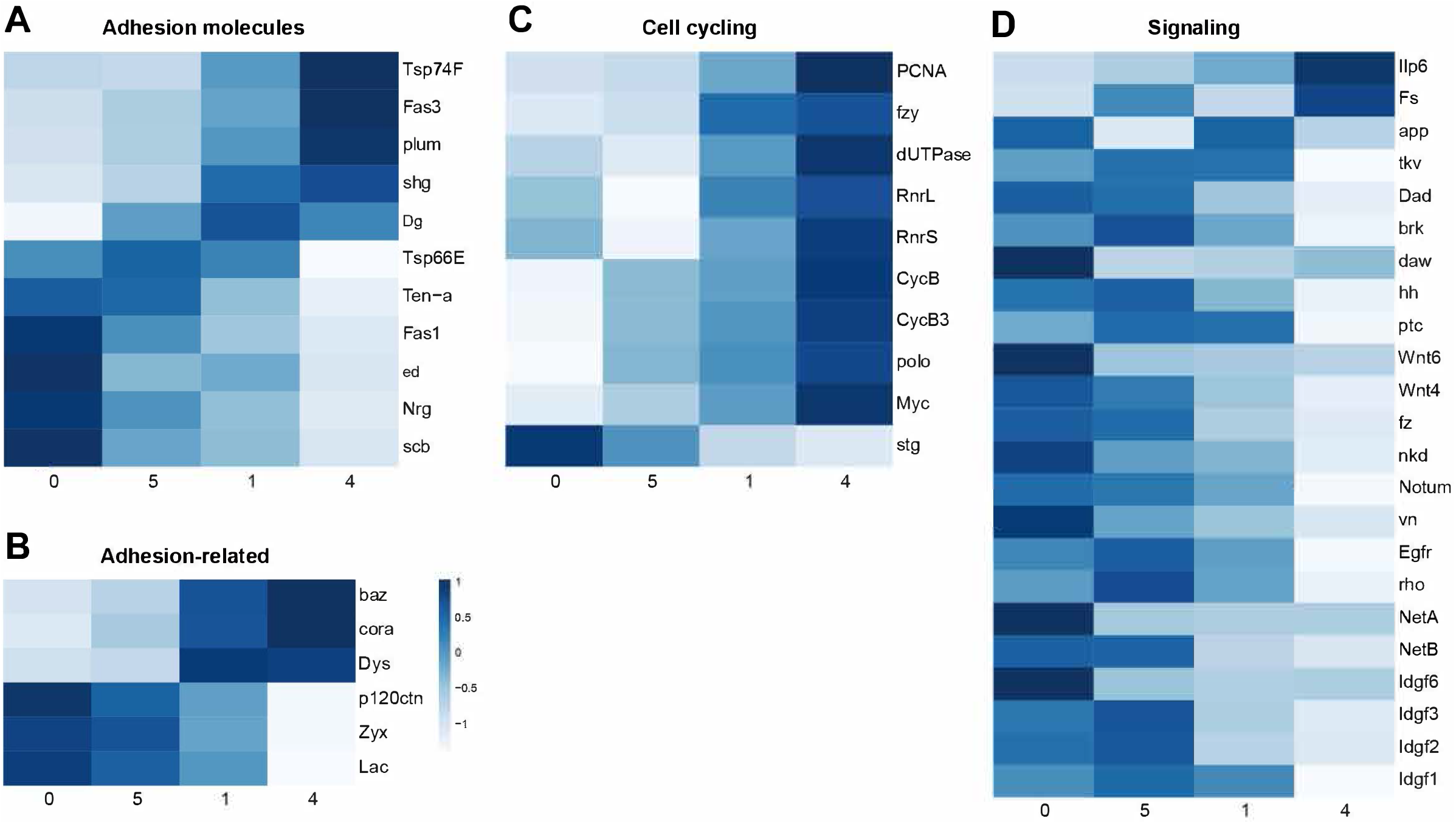
Potential regulators of cell behavior with AP-graded expression patterns. Heatmaps of selected genes encoding (A) adhesion molecules, (B) adhesion-related molecules, (C) drivers or regulators of cell cycling and (D) selected signaling molecules with pronounced differences in expression from ECS (groups 0 and 5), through FSCs (group 1) to early FCs (group 2).

Posterior FSC cycle faster than anterior FSCs, while ECs do not divide at all (13). Graded JAK-STAT activity, declining from the posterior, is one contributor to this patterned proliferation, while Hedgehog (Hh) and Phospho-inositide 3’ kinase (PI3K) pathways also provide major stimuli for FSC division (17, 44-46). Most ECs are found in G1, while most posterior FSCs are in G2, indicating limitation of cycling by G2/M restriction in all FSCs with an increasingly severe G1/S restriction further anterior (40). We found that several RNAs associated with DNA replication increased from anterior (group 0 and 5) to posterior (group 4), including *Proliferating Cell Nuclear Antigen* (*PCNA*), *deoxyuridine triphosphatase* (*dUTPase*), *Ribonucleotide reductase small* (*RnrS*) and *large* (*RnrL*) subunit genes. A similar, graded increase in drivers of mitosis was seen, including *fizzy* (*fzy*), *polo, cyclin B* (*cycB*) and *cyclin B3* (*cycB3*), though *string cdc25 phosphatase* (*stg*) had strongly graded expression of opposite polarity (Fig. 4C).

Signaling pathway activities can be indicated by expression of ligands, although their range of movement can be large, by receptors, which are sometimes up-regulated or down-regulated in response to signal and by genes that commonly respond transcriptionally to pathway activity, though often in tissue-specific ways. *hh* RNA was high in groups 0, 5 and 1, consistent with prior detection in anterior germarial cells, supplementing very strong expression in cap cells and terminal filament cells (47, 48). Correspondingly, the universal Hh target gene, *patched* (*ptc*) declines posteriorly from group 1 cell levels (Fig. 4D). *Wnt6* is expressed largely in group 0, consistent with previous evidence of expression only in anterior ECs, while *Wnt4*, which is strongly expressed in all ECs and FSCs, declines from group 0 through 5, 1 and 4 (36, 42, 48). A similar gradient is seen for *notum* and *naked* (*nkd*), which are sometimes positive transcriptional targets of Wnt signaling (49, 50). Those patterns are consistent with the observed anterior to posterior decline of a Wnt pathway reporter over the FSC domain (13, 51). *Epidermal Growth Factor Receptor* (*EGFR*), one of its ligands *vein* (*vn*), and *rhomboid* (*rho*), which processes Spitz EGFR ligand, all decline from group 5 through group 2 (Fig. 4D). Two genes often induced by BMP signaling, *brinker* (*brk*) and *Daughters against dpp* (*Dad*) show a similar decline. BMP signals supplied to GSCs are strictly limited to the anterior of the germarium (52). Activin signaling in the germarium has not been studied extensively but the agonist ligand *dawdle* (*daw*) was preferentially expressed anteriorly, with the converse pattern for the antagonist *Follistatin* (*Fs*) (53). The only other noted ligand with a strong posterior bias was an insulin receptor agonist, *Insulin-like peptide 6* (*Ilp6*). Expression of a set of imaginal disc growth factors (*Imaginal disc growth factor* (*Idgf*) 1, *2, 3, 6*) declined from anterior to posterior (Fig. 4D).

### Prospective FSC labeling for lineage analyses

The single-candidate method used to identify FSC locations has not commonly been used for other stem cells. Instead, for mouse stem cells the traditional approach has been to find a sufficiently specific marker to label a subset of cells through a recombination event and then determine by lineage analysis if such labeled cells have stem cell properties (2, 54). In practice, there is often no single suitable marker with absolute specificity. Instead, an empirical threshold of marker gene expression is engineered to initiate a *Cre-loxP* recombination event under specific conditions of sensitivity. Consequently, questions inevitably remain about whether all cells captured by that strategy are stem cells and whether only a subset of stem cells have been captured (2). These limitations are not commonly voiced but are significant.

ScRNA sequencing of germarial cells can provide candidate genes for targeting FSCs. The most common targeting strategy in Drosophila uses specific gene regulatory elements to drive yeast GAL4. GAL4 then activates a *UAS-flp* recombinase gene to initiate stable GFP expression in a recombinant cell and its derivatives (“G-trace”) (55). Temperature-sensitive GAL80 is generally included, so that the timing of recombination events can be limited to a chosen time window at the restrictive temperature. The activity of a specific *GAL4* transgene does not necessarily reflect normal expression of the parent gene and is normally assayed by including a *UAS-RFP* reporter. According to such RFP patterns, specific *Wnt4-GAL4, C587-GAL4* and *fax-GAL4* lines have been reported to be expressed largely or entirely anterior to the strong Fas3 border (56). We found that after at least 3d at the restrictive temperature, RFP expression was mostly anterior to the strong Fas3 border for all three lines, with strongest expression for *Wnt4-GAL4* and weakest for *fax-GAL4* (Fig. 5A-F). All *Wnt-GAL4* germaria (24/24) included RFP-positive cells anterior to the Fas3 border in locations we have previously designated as layer 1-3 FSCs, compared to 19/27 (70%) for *fax-GAL4*. Most such cells were labeled in each *Wnt-GAL4* germarium but only an average of 2.3 (44 total) for *fax-GAL4*. In about a third (8/24) of germaria for *Wnt4-GAL4* there was also detectable RFP in cells immediately posterior to the Fas3 border (generally one cell with weak expression) but RFP label was detected posterior to the Fas3 border in very few (3/59) *fax-GAL4* germaria. For *C587-GAL4*, RFP was detected after 7d at 30C in both layer 1-3 locations anterior to the Fas3 border and at least one cell posterior to the border in all cases (10/10). In the same samples, GFP patterns roughly mirrored RFP expression. However, while RFP expression provides an analog measure of GAL4 activity, GFP expression is digital, with permanent expression at a fixed level (driven by a *ubiquitin* promoter) once expression is triggered by a recombination event. Moreover, the threshold for triggering a stochastic recombination event can be quite low compared to generating a clear RFP signal. Hence, for lineage studies it is imperative to ascertain the initial GFP labeling pattern and not sufficient to rely on RFP expression patterns.

**Figure 5.**
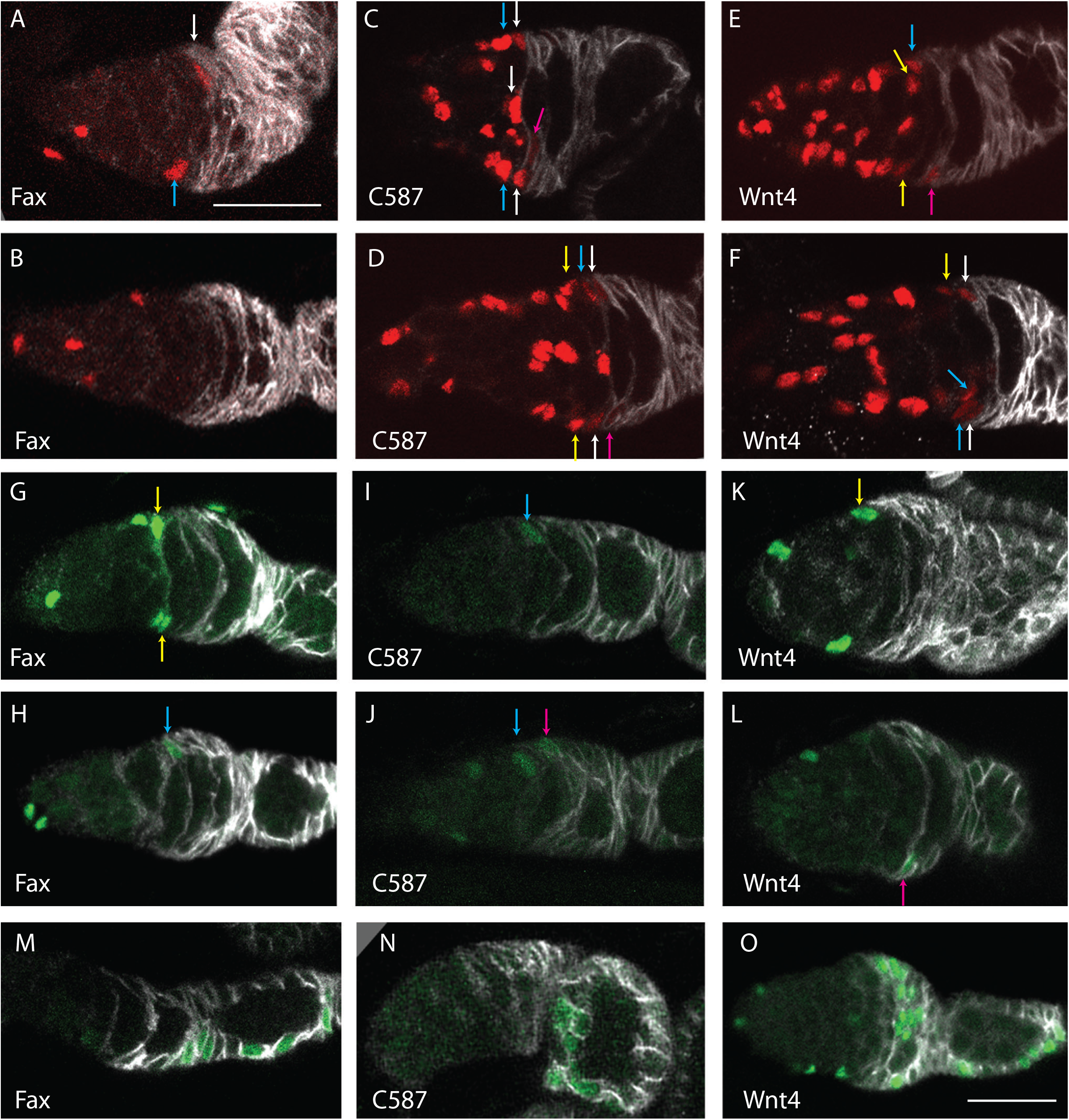
G-trace lineage analysis with *fax-GAL4, C587-GAL4* and *Wnt-GAL4*. (A-F) Expression of *UAS-RFP* after incubation at the restrictive temperature for (A, B, E, F) 3d or (C, D) 7d, with Fas3 antibody staining (white). In all cases, several ECs, far anterior (left) to strong Fas3 and arrows, express RFP (red). (A, B) RFP was seen in some (A) layer 1 (white arrow) and layer2 (blue arrow) cells but (B) not in all samples for *fax-GAL4*. (C, D) RFP was seen in layer 1 (white arrows), layer 2 (blue arrows) and layer 3 (yellow arrows), as well as more faintly in cells just posterior to the strong Fas3 border (pink arrows) for *C587-GAL4*. (E, F) RFP was seen in layer 1 (white arrows), layer 2 (blue arrows) and layer 3 (yellow arrows), as well as more faintly in cells just posterior to the strong Fas3 border (pink arrows) in (E) some samples but (F) not others for *Wnt4-GAL4*. (G-L) GFP (green) expression at d1 after incubation at the restrictive temperature (30C) and return to 18C. (G, H) For *fax-GAL4*, GFP was common in ECs, occasionally present in layer 1, 2 (blue arrow) or 3 (yellow arrows) cells (24%) and rarely present posterior to the strong Fas3 (white) border (6%). (I, J) For *C587-GAL4*, GFP was detected in ECs, sometimes in layer 1, 2 (blue arrows) or 3 cells (24%) and less often (J) immediately posterior to the strong Fas3 border (pink arrow) (14%). (K, L) For Wnt4-GAL4, GFP was detected in ECs, frequently in layer 1, 2 or 3 (yellow arrow) cells (56%) and almost as often (L) immediately posterior to the strong Fas3 border (pink arrow) (41%). (M-O) GFP (green) expression after a further 6d at 18C was found in small groups of FCs in the germarium or first three egg chambers in a subset of samples for (M) *fax-GAL4* (21%), (N) *C587-GAL4* (24%) and (O) *Wnt4-GAL4* (37%). Scale bar of (A) applies also to (B-N); all are 20µm.

Previous G-trace tests used long periods at the restrictive temperature and reported that many ovarioles later included GFP-labeled FCs for *Wnt4-GAL4* and *C587-GAL4* (56). Those results are broadly consistent with FSCs being located anterior to the strong Fas3 border. However, a similar experiment using *fax-GAL4* was reported to produce very few ovarioles with labeled FCs under normal conditions in well-fed flies (56). This evidence was used to argue that FSCs lie posterior to the strong Fas3 border. We repeated those conditions and found substantially different results. GFP-labeled FCs were seen in 34% (17/50) of ovarioles; each of these ovarioles also included labeled cells in layer 1-3 (with an average of close to 3 labeled layer 1-3 cells per germarium), while only 9/50 included labeled cells immediately posterior to the Fas3 border. These results, in contrast to the previous report (56), are consistent with layer 1-3 cells as a source of FCs and do not appear to be consistent with the suggestion of long-lived FSCs located posterior to the AP border. Under the same conditions, *C587-GAL4* (10/10= 100%) and *Wnt-GAL4* (24/24= 100% even by 3d) produced a greater frequency and density of FC labeling, but these two GAL4 lines are expressed at higher levels and in more cells anterior to the Fas3 border than *fax-GAL4*.

A much better lineage test would be to label single or very few cells in each germaria over a short time period, ensure no further labeling, and then “chase” for a period sufficient to determine if labeled FCs are formed. We experimented with a variety of conditions to achieve this outcome. A significant practical limitation here (and elsewhere (41, 57)) was the capacity of the temperature-sensitive *GAL80* transgenes to silence GAL4 activity at the permissive temperature. Even in the presence of two such transgenes, GFP clones were occasionally initiated for both *Wnt4-GAL4* and *fax-GAL4* for flies raised and aged at 18C. The nature of those GFP labeling patterns was variable but the frequent heavy-labeling of multiple ECs, FSCs and FCs suggested that most recombination events occurred during development (41), when the expression patterns of *Wnt4-GAL4* and *fax-GAL4* are potentially significantly different from those in adults. *C587-GAL4* with three copies of the *ts-GAL80* transgene gave virtually no background GFP-positive clones at 18C. A second experimental limitation concerns timing prior to determining initial GFP-labeling patterns. For the weakest GAL4 line, *fax-GAL4*, it was necessary to incubate for 10.5h at the restrictive temperature to have relevant cells labeled in at least 20% of ovarioles. Ideally, flies are then returned to 18C for long enough for all recombination events to manifest as detectable GFP signal before assaying a cohort of flies for initial GFP expression patterns. However, waiting too long will allow some initially labeled cells to divide or to move to different locations, including crossing the strong Fas3 boundary. We developed near-optimal timing for each GAL4 line but the compromise between a sufficient induction period and assaying a true time zero was greatest for *fax-GAL4* because it had the weakest expression. Conversely, the third ideal requirement of limiting GAL4 activity and initial GFP marking strictly to cells anterior to the strong Fas3 border was not met by *Wnt4-GAL4* and *C587-GAL4*. An ideal single GAL4 line, with strong and highly-selective expression, may not be attainable because of the inevitable similarities between a layer 1 FSC and the FC that it can become within a few hours.

Using the best conditions we could establish, we conducted quantitative G-trace experiments for all three GAL4 lines. Samples were scored to ascertain initial GFP labeling patterns (d1) and after a further 6d at 18C (d7). Since development is roughly twice as slow at 18C compared to 25C and egg chambers generally bud every 12h at 25C, it is expected that the most immature FCs (“immediate FCs”) will progress out of the germarium (2d) and occupy the 4th or 5th egg chamber, as previously observed directly for a 3d interval at 25C (9). A FSC may simply remain as a FSC but if it or its progeny become a FC, such FCs would be in the 4^th^ egg chamber or more anterior locations. Thus, based on our current understanding, we would expect that all marked FCs anterior to egg chamber 4 on d7 are derivatives of initially-marked FSCs (and a few FSC derivatives may also be present in the 4^th^ egg chamber).

For *Wnt4-GAL4*, a small number of ovarioles (<15%) at d1 and d7 included strong, widespread GFP labeling and presumably originated during development, as described above. These background clones were discounted and not included in totals because any additional labeling induced at the restrictive temperature in adults could not be scored. At d1, 19/34 germaria had cells labeled in layers 1-3 anterior to the strong Fas3 border (total of 37 such labeled cells; average 1.9), 14/34 had at least one GFP-positive cell immediately posterior to the Fas3 border (8 of these also had label in layers 1-3) (Fig. 5K, L). At d7, the observed fraction with marked FCs anterior to the 4^th^ egg chamber was 37% (11/30) (Fig. 5O). These results are compatible with the marked FCs at d7 originating from either layer 1-3 cells or cells posterior to the Fas3 border. Essentially, *Wnt-GAL4* initially labels cells posterior to the Fas3 border too frequently to provide a decisive test, even though the pattern of *UAS-RFP* labeling is strongly biased towards cells anterior to the Fas3 border.

For *C587-GAL4* at d1, 14/58 germaria had cells labeled in layers 1-3 anterior to the strong Fas3 border (total of 20 such labeled cells; average 1.4), 8/58 (14%) had a GFP cell immediately posterior to the Fas3 border (4 of these also had label in layers 1-3) (Fig. 5I, J). At 7d, 10/41 (24%) of ovarioles included labeled FCs anterior to the 4^th^ egg chamber and a further 5/41 (12%) included labeled cells in layers 1-3 with no labeled FCs (Fig. 5N). The proportion of 7d ovarioles with labeled FCs (24%) exceeded the proportion with labeled cells posterior to Fas3 on d1 (14%), suggesting that there must be a major source of FCs anterior to Fas3. By contrast, the proportion of germaria with labeled layer 1-3 cells at d1 and labeled FCs at d7 were similar, consistent with the former serving as FSCs.

For *fax-GAL4*, 15/123 ovarioles at d1 and 11/81 ovarioles at d7 showed heavily labeled background GFP signal and were excluded. At d1, 26/108 (24%) of germaria had cells labeled in layers 1-3 anterior to the strong Fas3 border (total of 70 such labeled cells; average 2.7), 7/108 (6%) had a GFP cell immediately posterior to the Fas3 border (6 of these also had label in layers 1-3) (Fig. 5G, H). At 7d, 15/70 (21%) of ovarioles included labeled FCs anterior to the 4^th^ egg chamber, FCs in the 4^th^ egg chamber were the only GFP-positive cells other than ECs in 4/70 (6%) ovarioles, and a further 4/70 (6%) included labeled cells in layers 1-3 with no labeled FCs (Fig. 5M). The proportion of 7d ovarioles with labeled FCs (24%) is much higher than the proportion with labeled cells posterior to Fas3 on d1 (6%), showing that the major source of FCs must be anterior to Fas3. The frequency of ovarioles with labeled layer 1-3 cells at d1 (24%) roughly matched the sum of FC derivatives (21%) and layer 1-3 cells with no FC derivative (6%) at d7, consistent with the expected behavior of layer 1-3 cells as FSCs (2, 13). FCs in the 4^th^ egg chamber (6%) may have derived from FSCs or cells immediately posterior to the Fas3 border (6%).

## Discussion

Single-cell RNA sequencing has great potential for understanding key regulators of a cell state. A first step in realizing this potential is to relate single-cell profiles to cell identities. For FSCs, the key intermediate is spatial location. Here, we used cells pre-sorted to represent mainly anterior germarial cells (ECs, FSCs and early FCs) and standard bioinformatic analyses to derive clusters guided by similarities. The clusters were notably similar to those of another study (37), facilitating assignment of group locations according to prior RNA *in situ* results for key indicators of those groups. The cluster profiles themselves showed a clear progression consistent with assignments along the AP axis and the placement of a key spatial landmark, the anterior border of strong surface Fas3 protein expression, roughly at the border between two groups (“1” and “4”). The anterior limit of group “1” was placed roughly three cells further anterior by comparison to RNA *in situs*, forming a cluster size of roughly the expected size to represent FSCs relative to the more anterior groups (“0” and “5”) representing ECs. Importantly, this study is the first to assign the FSC group by taking into account the most comprehensive functional investigation of FSC numbers and locations (13). Some other studies sampled the whole ovariole and therefore lacked sufficient FSCs in their samples (29, 30, 56); others did not acknowledge recent developments in understanding FSC biology or simply avoided an FSC designation, focusing instead on other cell types (37, 56, 57). Here, we included images of a variety of GFP-marked FSC lineages to illustrate FSC locations in 2-3 AP layers anterior to the Fas3 border, as determined previously by examining lineages with only a single candidate FSC (13). We also included a protocol for identifying the Fas3 border because other investigators have presented inconsistent and potentially confusing approaches and summaries (56).

We also undertook a complementary approach to verify FSC locations through prospective “G-trace” labeling (55). We obtained results consistent with our prior designation of FSCs occupying 2-3 layers anterior to the Fas3 border and contradicting an assertion that FSCs are posterior to the AP border (56), based principally on using the very same *fax-GAL4* reagent to initiate clones. Crucially, our tests involved carefully controlled conditions that allowed production and measurement of a suitable number of GFP-labeled cells during a short labeling period, followed by a chase period to ascertain the fate of such cells. Results with *fax-GAL4* and *C587-GAL4* to initiate labeling were consistent with FC-producing cells lying anterior to the Fas3 border, although that conclusion was less numerically robust for *C587-GAL4* because the associated initial GFP-labeling pattern was less specific than for *fax-GAL4*.

The assignment of a group of scRNA expression profiles to FSCs is not precise for a number of reasons, including imperfect and limited depth of RNA profiles, latitude in spatial correspondence according to RNA *in situs*, and the systematic limitation that FSCs become ECs or FCs within a few hours and therefore inevitably have strong similarities to those neighboring derivatives. Nevertheless, the results of this study have the unique virtue that the approximately-defined FSC group of cells is in accord with current understanding of FSC numbers and locations determined by thorough functional analyses (2, 9, 13, 17, 58-60). FSC behavior is guided by a number of external signals but with prominent roles for two graded signaling pathways, Wnt and JAK-STAT (13, 17). RNAs that also have graded AP expression from ECs through to early FCs are therefore of special interest as candidate mediators of spatially appropriate behavior guided by these graded signals. We have highlighted such RNAs encoding potential effectors of cell movements and differentiation or cell division as candidates for further investigation of functional roles and potential transcriptional regulation by Wnt or JAK-STAT pathways.

## Supporting information

Supplementary Material Description

## Acknowledgments

This work was supported by the National Science Foundation of China 32172467 to JH, NIH RO1 GM079351 to DK, and a SURF undergraduate research fellowship to XT. We thank John Lis (Cornell University) and the Bloomington stock center for provision of genetic reagents, the Developmental Studies Hybridoma Bank (DSHB) for antibodies, FlyBase as an information resource, David Melamed for preparation of Figure 4, and the confocal microscope resource provided by the Dept. of Biological Sciences, Columbia University.

## Materials and Methods

### Single-cell Suspension Preparation and Fluorescence-activated Cell Sorting (FACS)

Female flies of genotype *C587-GAL4* / *yw; UAS-CD8-RFP*/*+* were selected on the day of eclosion and maintained on rich food at 25C together with males for 4-7d. More than 200 pairs of ovaries were dissected in ice cold DPBS (Dulbecco’s Phosphate Buffered Saline) solution, and then gently washed twice by DPBS. The ovaries were digested in 700 µl solution containing 0.5% trypsin (Sigma, #T4799) and 0.25% collagenase (Invitrogen, #17018-029) at 25C for about 20 min with gentle shaking. The dissociated cells were filtered through a 40-micron cell strainer (Falcon, #352340), and then Centrifuged at 400 g at 4C for 5 min. Supernatant was removed and the cell pellet was resuspended in DPBS with 0.2% BSA. Cell suspensions were sorted using a FACS Aria Ⅲ sorter MoFlo XDP (Beckman-Coulter) based on RFP signal. Cells from w^1118^ fly ovaries were used to set the negative fluorescence gate for the RFP panel. RFP-positive cells were sorted into DPBS containing 0.2% BSA, pelleted (400 g for 10 min) and resuspended in DPBS. The LUNA-FL double fluorescent cell counter (Logos Biosystems) was used to count cell number and the live/dead cell ratio. The suspension of single cells, with > 85% live cells, had a final concentration of 500 cells/µl.

### Single cell RNA-sequencing (scRNA-seq)

The sorted cells were loaded on a Chromium Single cell Controller (10 X Genomics) using the Chromium Single-cell 3’ Library & Gel Bead kit v3 according to manufacturer’s directions. Briefly, the Gel Beads containing the poly-T primer sequence were linked with a cell barcode and UMI (unique molecular identifier) and the cells are wrapped by “oil droplets” to form a GEM (Gel Bead in Emulsion). Cells were lysed in these droplets and reverse-transcribed to form full-length cDNA sequence. After the oil droplets were broken and purified, the cDNA library was PCR amplified and ligated with sequencing primers. cDNA library quantification assays and quality check analysis were performed using an Agilent 2100 Bioanalyzer and Thermo Fisher Qubit Fluorometer. The library samples were then diluted to a 10 nM concentration and sequenced on an Illumina NovaSeq 6000 platform (Illumina, San Diego) for PE150 sequencing by Annoroad Gene Tech. Co., Ltd (Beijing, China). A total of 377, 489, 707 reads were obtained for the sample, with 290, 154 mean reads per cell.

### Raw data preprocessing

The raw sequencing data was processed with alignment, barcode assignment and UMI counting by Cell Ranger (version 6.1.1) using the previously published pipeline (61). The reference index for Cell Ranger was built using the Drosophila melanogaster genome (version BDGP6.32) available on the Ensembl genome database.

### Single cell RNA-seq data analysis

Cell Ranger’s raw gene count matrix was further analyzed using the Seurat (v4.0.2) R package with standard protocols (62). Only genes expressed in more than 3 cells were kept, and only cells that have unique feature counts between 100 and 4,500, and less than 20% mitochondrial counts were selected for further analysis. Gene expression results were log-normalized, and then regressed on the percentage mitochondrial gene content. The top 16 principal components of Principal component analysis (PCA) were used for clustering according to the Elbow Plot (resolution = 0.9). The ECs, FSC and FCs were identified by markers of each cluster. tSNE plots were used to visualize the clustering results. Mean expression values for each cluster were calculated by “AverageExpression” in Seurat. The R package “pheatmap” was used to generate heatmaps and the gene expression levels were scaled by row.

### MARCM and multicolor FSC lineages

In a typical MARCM (mosaic analysis with repressible cell marker) experiment, 1-3d old adult *Drosophila melanogaster* females with the appropriate genotype (*yw hs-Flp, UAS-nGFP, tub-GAL4* /*yw; act-GAL80 FRT40A* / *FRT40A; act>CD2>GAL4*/ *+* or *yw hs-Flp, UAS-nGFP, tub-GAL4* /*yw; FRT42D act-GAL80 tub-GAL80* / *FRT42D; act>CD2>GAL4*/ *+* were given a single 30 min heat shock at 37C. Afterwards, flies were incubated at 25C and maintained by frequent passage on normal rich food supplemented by fresh wet yeast before dissection 6 or 12d later. For multicolor lineages the procedure was similar but with multiple heat shocks (13) and the final genotype of *yw hs-flp* / *yw; tub-lacZ FRT40A FRT42B* / *ubi-GFP FRT40A FRT42B His2Av-mRFP*. Adult female ovaries were dissected in PBS, leaving the tip of the abdomen attached, and fixed in 4% paraformaldehyde (Electron Microscopy Sciences, #15710) for 10 minutes at room temperature, covered to prevent bleaching of fluorophores. The ovaries were then washed three times with 1X PBST (PBS with 0.1% Triton X-100 and 0.05% Tween 20) and blocked at room temperature in blocking solution (PBST with 10% normal goat serum) for 30 minutes. Primary antibodies mouse anti-Fasciclin 3 (Developmental Studies Hybridoma Bank, #7G10, 1:250), and rabbit anti-GFP (Invitrogen, #A6455, 1:1000) were added and samples were nutated for an hour at room temperature. The ovaries were then rinsed with 1X PBST three times for 10 minutes each before being incubated with secondary antibodies Alexa Fluor-647 goat anti-mouse (Invitrogen, #A21236, 1:1000) and Alexa Fluor-488 goat anti-Rabbit (Invitrogen, #A11034, 1:1000) at room temperature for 1 hour. The ovaries were then rinsed twice with PBST and once in PBS. Each ovary pair was then broken apart in PBS on a slide and mounted using *DAPI fluoromount-G* mounting medium (Southern BioTech, OB010020). Ovarioles were imaged with a Zeiss LSM700 or LSM800 confocal microscope, operated in part by the Zeiss ZEN software.

### Protocol for identifying cell locations relative to landmark of anterior border of strong Fas3 staining

A variety of indicators help to ensure reproducible determination of the anterior border of strong Fas3 staining. These include germarium width, identifying the youngest stage 2b cyst and noting cell process locations. Starting with a mid-z-section, identify the youngest stage 2b cyst by the criteria of spanning the germarium and not being at all rounded. Most germaria have only a single candidate 2b cyst with a clearly more rounded stage 3 cyst more posterior. In such germaria, the Fas3 border runs along the posterior surface of the 2b cyst. In other germaria (up to about a quarter), a new 2b cyst has formed as a more posterior 2b cyst just starts to round. Here, the strong Fas3 border lies between the two 2b cysts. The strong Fas3 border can then be followed as a continuous surface through neighboring z-sections. Layer 1 cells immediately anterior to the Fas3 border have strong Fas3 staining on their posterior surface but weak or incomplete outlining of the anterior surface by Fas3. Labeling in MARCM clones usually allows visualization of FSC processes (even faintly when using a nuclear-targeted GFP marker). Layer 1 cell processes are present along the Fas3 border, whereas layer 2 processes are anterior to the stage 2b cyst. The widest part of the germarium is generally very close to the Fas3 border, with layer 1 FSCs at the widest location in about three-quarters of samples and immediate FCs at that location in the rest.

### G-trace experiments

Flies of genotype *UAS-RFP, UAS-flp, ubi>stop>GFP* / *CyO; tub-GAL80(ts)* / *TM2* were crossed to (a) *tub-GAL80(ts)* / *CyO; fax-GAL4* / *TM2* or (b) *Wnt-GAL4* / *CyO; tub-GAL80(ts)* / *TM2* or (c) *C587-GAL4; tub-GAL80(ts)* / *CyO; tub-GAL80(ts)* / *TM2* flies at 18C to collect experimental progeny with no balancer chromosomes and two (for *fax-GAL4* and *Wnt4-GAL4*) or three (for *C587-GAL4*) copies of *tub-GAL80(ts)*. Young (1-4d old) experimental flies were then shifted to 29C or 30C for various periods and, in some cases, returned to 18C, as described in the Results for specific tests. For the final lineage tests the conditions were 3h at 30C followed by 13h at 18C (*Wnt-GAL4*), 6h at 30C followed by 13h at 18C (*C587GAL4*), or 10.5h at 30C followed by 13h at 18C (*fax-GAL4*) before dissecting for the first time-point, followed by a further 6d at 18C before dissecting another cohort. An equivalent cohort of flies was kept at 18C throughout. In all cases, flies were transferred to fresh food every 2-3d. Dissection, staining, mounting and analyses were as described for MARCM experiments.

## Supplementary Materials

**Figure S1. Identity markers for peripheral groups in initial t-SNE clusters**. Related to Figure 2. (A-C) t-SNE plots of complete data set (Fig. 2A) showing color-coded relative expression levels of genes characteristic of (A) TF or cap cells (*Lmx1a, en, dpp, wg*), (B) stalk or pre-stalk cells (*LamC, sim*) or (C) germline cells (*zpg, chinmo, ovo* and *otu*).

**Supplementary Table 1**

Ranked list of genes, ordered by adjusted p-value (p_val_adj), for each cluster (0, 5, 1, 4, 2, 3), showing the Log fold-change of the average expression between clusters and the percentage of cells where the gene is detected in the cluster. Clusters correspond to Fig. 2C.

**Supplementary Table 2**

Genes with *increasing* expression from cluster 0 through clusters 5 and 1 to cluster 4. These genes were picked by calculating the Spearman correlation coefficient to find genes with a positive coefficient greater than 0.4. Average expression value for each cluster (including clusters 2 and 3) is listed (“AverageExpression”, described in Methods). The change from cluster 0 to cluster 4 (“updegree”) is calculated as (cluster4 – cluster0)/cluster4. The resulting values were classified into quartiles, where the lowest number represents the greatest variation between groups 0 and 4. Genes are ordered according to the absolute expression value in cluster 4.

**Supplementary Table 3**

Genes with *decreasing* expression from cluster 0 through clusters 5 and 1 to cluster 4. These genes were picked by calculating the Spearman correlation coefficient to find genes with a negative coefficient greater than 0.4 magnitude. Average expression value for each cluster (including clusters 2 and 3) is listed (“AverageExpression”, described in Methods). The change from cluster 0 to cluster 4 (“updegree”) is calculated as (cluster0 – cluster4)/cluster4. The resulting values were classified into quartiles, where the lowest number represents the greatest variation between groups 0 and 4. Genes are ordered according to the absolute expression value in cluster 0.

