## Supplementary Figure 1 for "Determination of expression profiles for Drosophila ovarian Follicle Stem Cells (FSCs) using single-cell RNA sequencing"

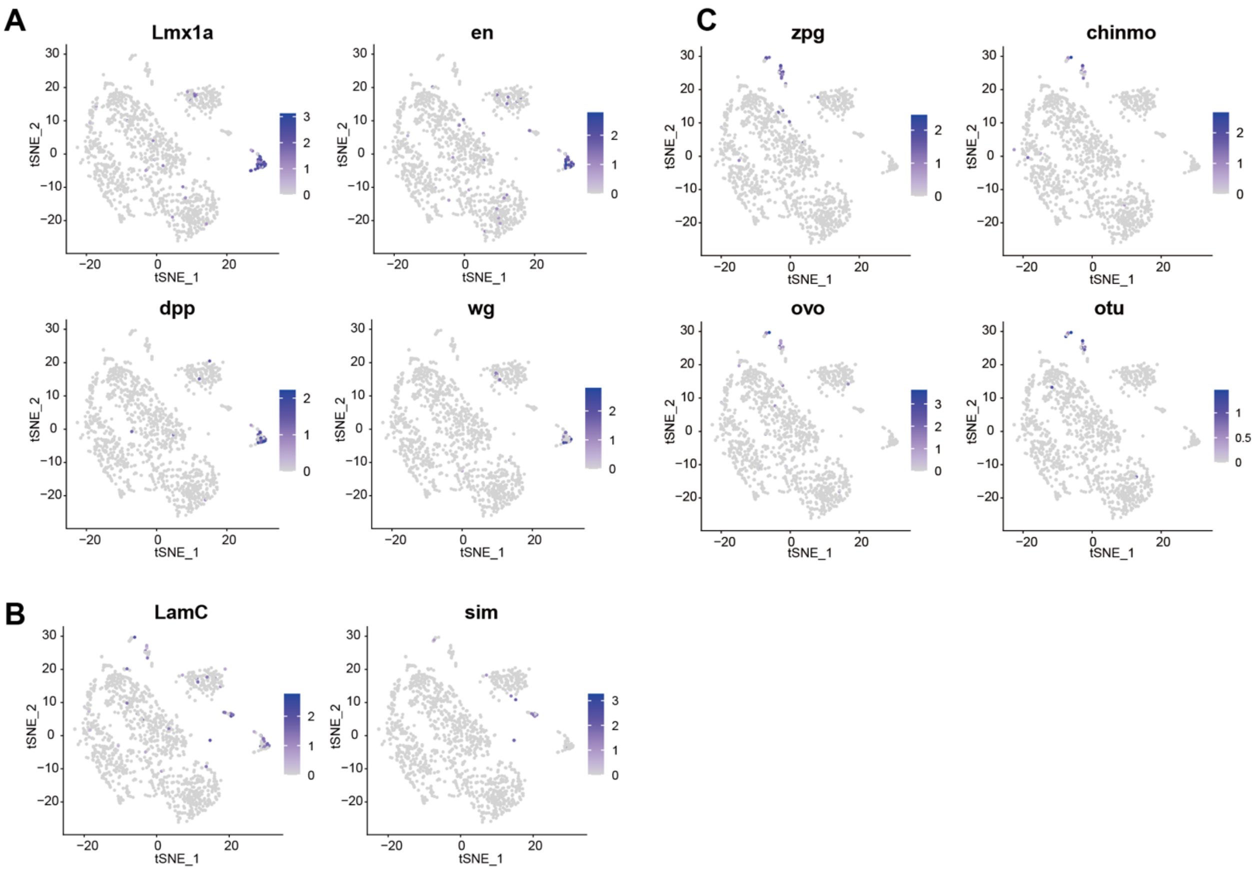


**Figure S1. Identity markers for peripheral groups in initial t-SNE clusters.** Related to Figure 2. (A-C) t-SNE plots of complete data set (Fig. 2A) showing color-coded relative expression levels of genes characteristic of (A) TF or cap cells (*Lmx1a*, *en*, *dpp*, *wg*), (B) stalk or pre-stalk cells (*LamC*, *sim*) or (C) germline cells (*zpg*, *chinmo*, *ovo* and *otu*).
